# Cation and anion channelrhodopsins: Sequence motifs and taxonomic distribution

**DOI:** 10.1101/2021.03.23.436664

**Authors:** Elena G. Govorunova, Oleg A. Sineshchekov, Hai Li, Yumei Wang, Leonid S. Brown, Alyssa Palmateer, Michael Melkonian, Shifeng Cheng, Eric Carpenter, Jordan Patterson, Gane K.-S. Wong, John L. Spudich

## Abstract

Cation and anion channelrhodopsins (CCRs and ACRs, respectively) primarily from two algal species, *Chlamydomonas reinhardtii* and *Guillardia theta*, have become widely used as optogenetic tools to control cell membrane potential with light. We mined algal and other protist polynucleotide sequencing projects and metagenomic samples to identify 75 channelrhodopsin homologs from three channelrhodopsin families, including one revealed in dinoflagellates in this study. We carried out electrophysiological analysis of 33 natural channelrhodopsin variants from different phylogenetic lineages and 10 metagenomic homologs in search of sequence determinants of ion selectivity, photocurrent desensitization, and spectral tuning in channelrhodopsins. Our results show that association of a reduced number of glutamates near the conductance path with anion selectivity depends on a wider protein context, because prasinophyte homologs with the identical glutamate pattern as in cryptophyte ACRs are cation-selective. Desensitization is also broadly context-dependent, as in one branch of stramenopile ACRs and their metagenomic homologs its extent roughly correlates with phylogenetic relationship of their sequences. Regarding spectral tuning, two prasinophyte CCRs exhibit red-shifted spectra to 585 nm, although their retinal-binding pockets do not match those of previously known similarly red-shifted channelrhodopsins. In cryptophyte ACRs we identified three specific residue positions in the retinal-binding pocket that define the wavelength of their spectral maxima. Lastly, we found that dinoflagellate rhodopsins with a TCP motif in the third transmembrane helix and a metagenomic homolog exhibit channel activity.

**IMPORTANCE:** Channelrhodopsins are widely used in neuroscience and cardiology as research tools and are considered as prospective therapeutics, but their natural diversity and mechanisms remain poorly characterized. Genomic and metagenomic sequencing projects are producing an ever-increasing wealth of data, whereas biophysical characterization of the encoded proteins lags behind. In this study we used manual and automated patch clamp recording of representative members of four channelrhodopsin families including a family that we report in this study in dinoflagellates. Our results contribute to a better understanding of molecular determinants of ionic selectivity, photocurrent desensitization, and spectral tuning in channelrhodopsins.

## INTRODUCTION

Channelrhodopsins (ChRs) are light-gated ion channels initially discovered in chlorophyte algae, in which they serve as photoreceptors guiding phototactic orientation (1–3). Subsequently, ChRs have also been found in the genomes/transcriptomes of cryptophyte and haptophyte algae (4, 5), the heterotrophic protists known as Labyrinthulea (5) and giant viruses that infect marine microorganisms (6, 7). Ongoing polynucleotide sequencing projects provide a rich hunting ground for further exploration of ChR diversity and taxonomic distribution.

Functionally, ChRs are divided into cation and anion channelrhodopsins (CCRs and ACRs, respectively) (8). Both ChR classes serve for photocontrol of excitable cells, such as neurons and cardiomyocytes, via a biotechnique known as optogenetics (9, 10). However, structural determinants for cation and anion selectivity in ChRs remain poorly understood. X-ray crystal structures (11–15) indicate that the ion conductance path in algal ChRs is formed by transmembrane helices (TM) 1, 2, 3 and 7. All so far known ACRs contain a non-carboxylate residue in the position of the protonated Schiff base counterion in bacteriorhodopsin (Asp85), whereas in nearly all CCRs this carboxylate is conserved. However, this sequence feature cannot be regarded as a sole indicator of anion selectivity, because some chlorophyte CCRs also show a non-carboxylate residue in the counterion position (e.g. *Ds*ChRl from *Dunaliella salina* (16)).

Most chlorophyte CCRs contain five Glu residues in TM2 and the TM2-TM3 loop (Glu82, Glu83, Glu90, Glu97, and Glu101 in ChR2 from *Chlamydomonas reinhardtii*, *Cr*ChR2), whereas in all so far known ACRs most or even all of the corresponding positions are occupied with non-carboxylate residues. Therefore, it has been proposed that negative electrostatic potential of the channel pore defines cation selectivity (17, 18). Indeed, mutagenetic remodeling of the pore to reduce electronegativity yielded permeability for anions in chlorophyte CCRs (17–21). Yet, some of the TM2 glutamates are conserved in ACRs and apparently do not interfere with their anion conductance.

Other biophysical properties of ChRs relevant for optogenetic applications are their desensitization under continuous or pulsed illumination (also called “inactivation” in the literature) and spectral sensitivity. In an earlier study, a group of ACRs discovered in the TARA marine transcriptomes demonstrated particularly rapid and strong desensitization (22). As their source organisms were not known, these proteins were named MerMAIDs (<u>Me</u>tagenomically discove<u>r</u>ed, <u>M</u>arine, <u>A</u>nion-conducting and <u>I</u>ntensely <u>D</u>esensitizing channelrhodopsins). However, strong desensitization cannot serve as a characteristic of a single ChR family, because it was also observed in some “bacteriorhodopsin-like” CCRs (BCCRs) from cryptophytes that show very little sequence homology with MerMAIDs (23).

To gain more insight into the taxonomic distribution and structure-function relationships of ChRs, we identified 75 ChR homologs from several phylogenetic lineages and metagenomic samples, and tested 27 of them along with 16 previously reported sequences by heterologous expression in cultured mammalian cells followed by patch clamp recording. We show that the same pattern of conserved Glu residues may accompany cation or anion conductance in ChRs from different taxa, and that the degree of desensitization in MerMAID homologs is the greater, the closer their sequences are to those of the first reported MerMAIDs. We report two prasinophyte CCRs with red-shifted spectra and confirm that three specific residues in the retinal-binding pocket are responsible for wavelength regulation in cryptophyte ACRs. Finally, we demonstrate that some dinoflagellate rhodopsins possess channel activity.

## RESULTS

### Prasinophyte CCRs

Only a few of >150 so far identified chlorophyte ChRs (Fig. 1, and Data Sets S1 and S2) have been tested by heterologous expression. Both *C. reinhardtii* ChRs conduct cations (2, 3), so other chlorophyte ChRs were also assumed to be CCRs. However, a recent study has demonstrated that two ChRs from the prasinophyte genus *Pyramimonas* in fact conduct anions (6), which called for a more detailed functional analysis of chlorophyte ChRs.

**Figure 1.**
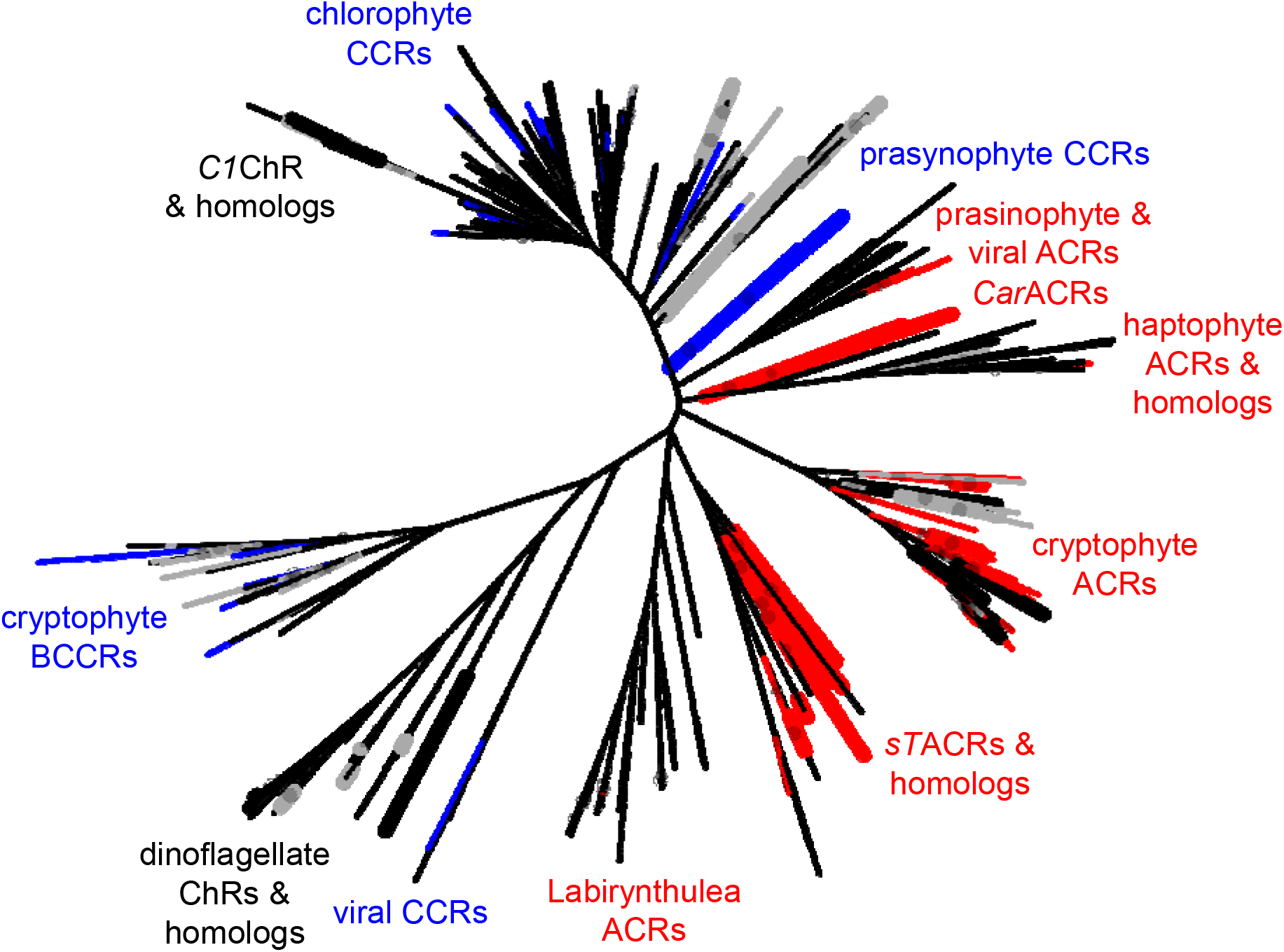
An unrooted phylogenetic tree of ChRs. The nodes are color-coded as following: red, confirmed anion selectivity; blue, confirmed cation selectivity; gray, non-functional; black, ion selectivity not determined. Thicker nodes show ChRs characterized in this study. Gray circles show ultrafast bootstrap support values above 95%. A tree file in the Newick format is available as Data Set S2, and the corresponding protein alignment, as Data Set S3.

Three ChR homologs derived from the prasinophytes *Crustomastix stigmatica*, *Mantoniella squamata* and *Pyramimonas melkonianii* (6) exhibit a residue pattern typical of cryptophyte ACRs, i.e. they display conserved Glu82 and Glu90 with non-carboxylate residues in the positions of Glu83, Glu97, Glu101 and Glu123 of *Cr*ChR2 (Fig. 2A). In a *Cymbomonas tetramitiformis* sequence (6), Glu90 and Glu97 are conserved, whereas Glu82 is replaced with Gln (Fig. 2A). We synthesized mammalian codon-adapted versions of these rhodopsin domains, fused them with C-terminal enhanced yellow fluorescent protein (EYFP), expressed in HEK293 (human embryonic kidney) cells and analyzed by manual whole-cell patch clamp.

**Figure 2.**
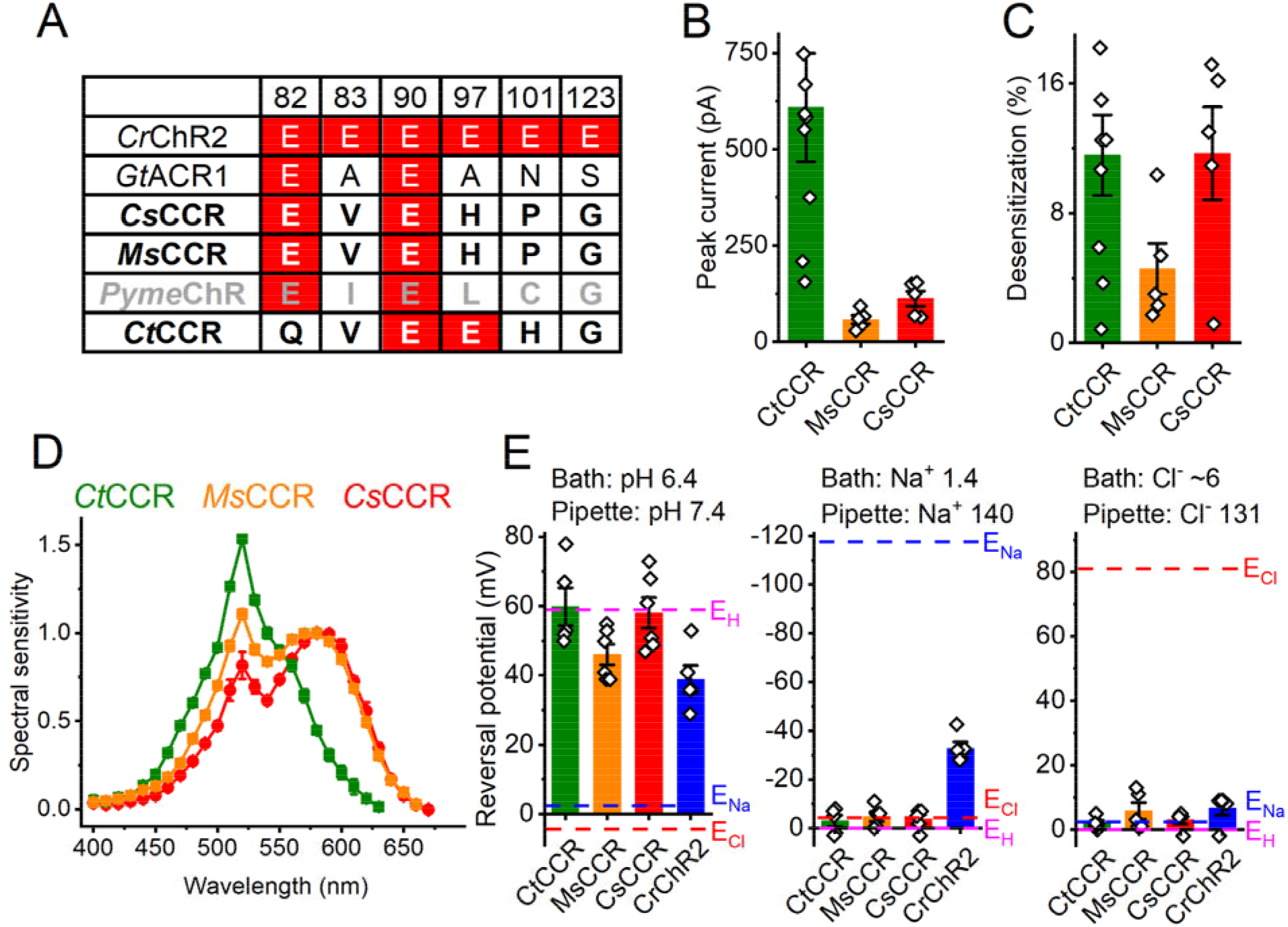
Prasinophyte CCRs. (A) Amino acid residues corresponding to the indicated positions in *Cr*ChR2. ChRs characterized in this study are in bold (black, functional; gray, non-functional). Conserved glutamates are highlighted red. (B) Peak photocurrent amplitudes generated at −60 mV in response to 1-s light pulses at the wavelength of the spectral maximum. (C) Desensitization of photocurrents after 1-s illumination. (D) Action spectra of photocurrents. The data points show mean ± sem (n = 4-8 scans). (E) Reversal potentials of photocurrents. The bars in B, C and E show mean ± sem; diamonds, data from individual cells.

Three of these ChRs generated photocurrents (Fig. 2B and C) in our standard buffer system (for solution compositions see Table S1), whereas the homolog from *P. melkonianii* that we named *Pyme*ChR was non-electrogenic. The action spectra of photocurrents generated by the *M. squamata* and *C. stigmatica* homologs were red-shifted (the rhodopsin maxima at ~580 and 585 nm, respectively; Fig. 2D). Their retinal-binding pockets are nearly identical, but differ from those of previously known red-shifted ChRs (Fig. S1A). Both spectra exhibited a second band at ~520 nm that reflected a Förster resonance energy transfer (FRET) from EYFP to rhodopsin, as was earlier shown in RubyACRs from Labyrinthulea (5). The efficiency of FRET was even greater in the homolog from *C. stigmatica*, the rhodopsin peak of which was observed in the green spectral region and could not be accurately resolved because of the FRET contribution (Fig. 2D, olive line).

To test the relative permeability of the prasinophyte homologs for H^+^, Na^+^ and Cl^−^ we varied the concentration of each of these ions in the bath (for solution compositions see Table S1), measured the current-voltage relationships and determined the reversal potentials (E_rev_). *Cr*ChR2 was included in this experiment for comparison. Fig. 2E shows that under all tested conditions E_rev_ for all three homologs was close to the equilibrium potential of H^+^, indicating that they are H^+^-selective channels with negligible permeability for Na^+^ and Cl^−^. We named them *Ct*CCR, *Ms*CCR, and *Cs*CCR. A less positive E_rev_ of *Ms*CCR photocurrents probed under the H^+^ gradient does not result from the permeability for Na^+^ as it does in *Cr*ChR2, and most likely reflects a contribution of intramolecular charge transfers, as previously found in other CCRs (24).

Four sequences from Chlorophyceae have only the Glu82 homolog, as do prasynophyte ACRs (Fig. S1B), but show no close sequence homology to them. Four sequences from Chlorodendrophyceae contain no glutamate residues in any of the six analyzed positions (Fig. S1D) and form a separate branch on the phylogenetic tree (Fig. 1). Very unusually, in the latter sequence group the Asp residue corresponding to Asp212 of bacteriorhodopsin is located not four residues upstream, as in most known microbial rhodopsins, but three residues upstream of the Schiff base lysine (Fig. S1E). None of these eight proteins, nor a ChR homolog from the streptophyte *Coleochaete* generated channel currents (see Supplemental Text and Fig. S1C for details).

### Stramenopile ACRs and their metagenomic homologs

The first MerMAIDs reported were seven homologous ACRs identified in metagenomic samples (22). Recently, close homologs were found in unclassified stramenopile species (5, 25), which suggests that the original MerMAIDs also originate from stramenopiles. We have identified 20 additional MerMAID homologs, nine haptophyte ACR homologs and two Labyrinthulea ACR homologs (Data Set S1) in metagenomic databases (Data Set S4). We tested EYFP fusions of five <u>m</u>eta<u>g</u>enomic MerMAID homologs (abbreviated by the lower case “mg” in the protein name), two closely related sequences from the unclassified <u>s</u>tramenopile strain <u>T</u>OSAG23-3 (abbreviated by “*sT*” (5)), and three sequences from the bicosoecid stramenopile *<u>Ca</u>feteria <u>r</u>oenbergensis* (abbreviated by “*Car*” to distinguish them from *C. reinhardtii* ChRs). In most sequences of this group both Glu82 and Glu90 (CrChR2 numbering) are conserved, as in the previously known cryptophyte ACRs and MerMAIDs (Fig. 3A). The five tested MerMAID homologs and those from TOSAG23-3 clustered together with the first reported MerMAIDs (Fig. S2), whereas Cafeteria homologs formed a separate branch related to haptophyte ACRs (Fig. 1). Each of these homologs generated photocurrents in HEK293 cells (Fig. 3B). As shown below, these ChRs conduct anions, so we designated them ACRs.

**Figure 3.**
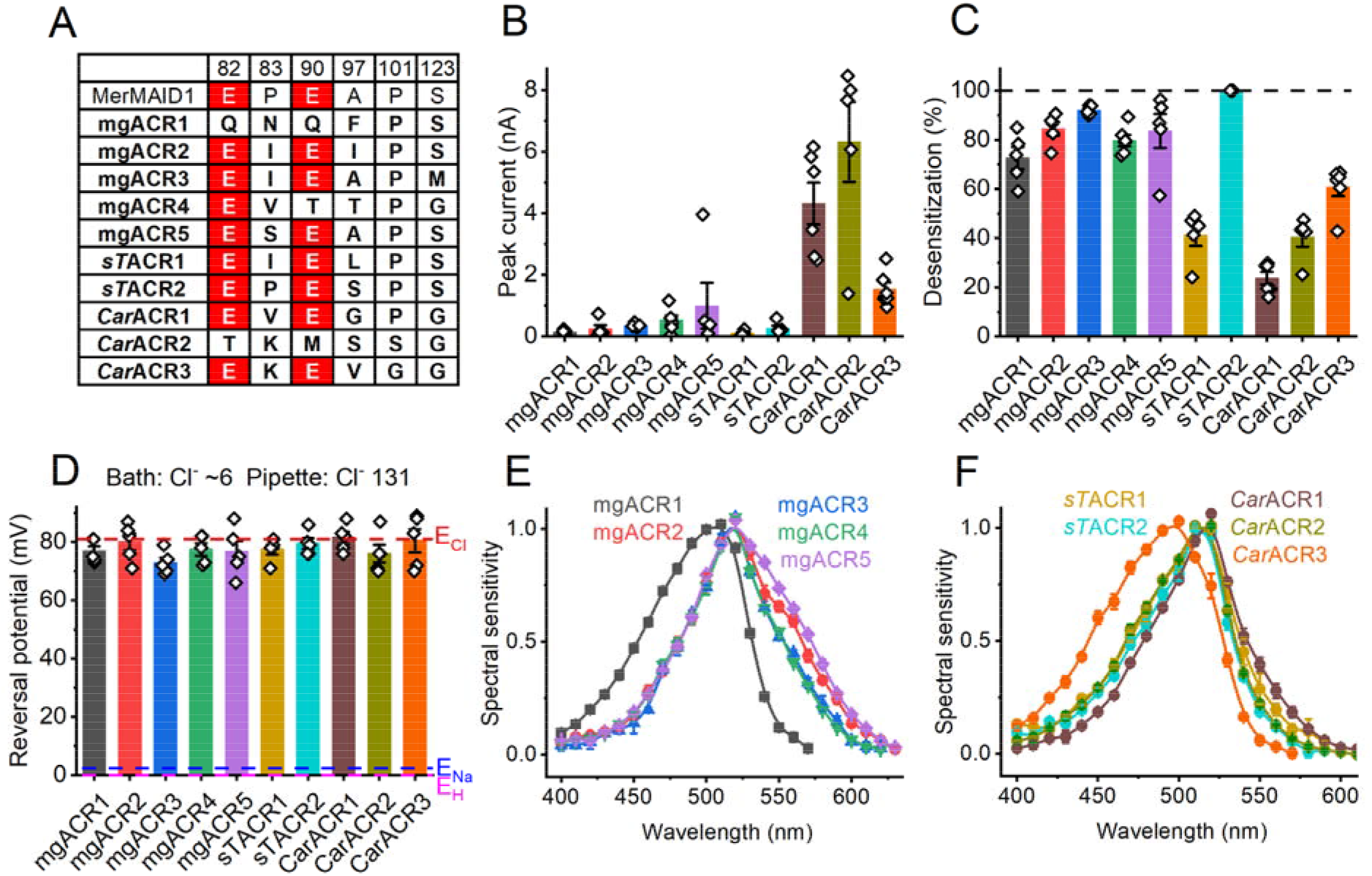
Stramenopile ACRs and their metagenomic homologs. (A) Amino acid residues in the ion conductance pathway, corresponding to the indicated positions in *Cr*ChR2. ChRs characterized in this study are in bold face. Conserved glutamates are highlighted red. (B) Peak photocurrent amplitudes generated at −60 mV in response to 1-s light pulses at the wavelength of the spectral maximum. (C) Desensitization of photocurrents after 1-s illumination. (D) Reversal potentials of photocurrents. In B-D the bars show mean ± sem; empty diamonds, data from individual cells. (E and F) Action spectra of photocurrents. The data points are mean ± sem (n = 4-6 scans).

Out of all tested homologs, *s*TACR2 is the most closely related to the first reported MerMAIDs, which exhibit nearly complete desensitization (Fig. S2). Similarly, *s*TACR2 photocurrents showed nearly complete desensitization (Fig. 3C, cyan), whereas photocurrents from *s*TACR1, the most distant homolog (Fig. S2), exhibited only ~40% desensitization (Fig. 3C, dark yellow). The values for other homologs were intermediate (Fig. 3C). All ChRs of this group demonstrated exclusively anion permeability (Fig. 3D). The action spectra of their photocurrents are shown in Fig. 3E and F. The shape of some spectra (e.g. mgACR2 and mgACR5) indicated a contribution of FRET from EYFP.

### Cryptophyte ACRs

Cryptophytes are the taxon in which the first natural ACRs were discovered (4). To explore the diversity of cryptophyte ACRs further, we have analyzed transcriptomes of 20 additional cryptophyte strains (Table S2) and identified 15 transcripts homologous to previously known cryptophyte ACRs (Data Set S1). As no species names have been assigned to their source organisms, we used the numbers 3-8 in the abbreviated protein names to designate different Rhodomonas strains (the numbers 1 and 2 have already been assigned to the previously analyzed strains). The Glu82 homologs is conserved in all, and the Glu90 homolog, in most of these proteins, whereas all other analyzed positions are occupied with non-carboxylate residues (Fig. 4A). Thirteen homologs generated photocurrents upon expression in HEK293 cells (Fig. 4B and C).

**Figure 4.**
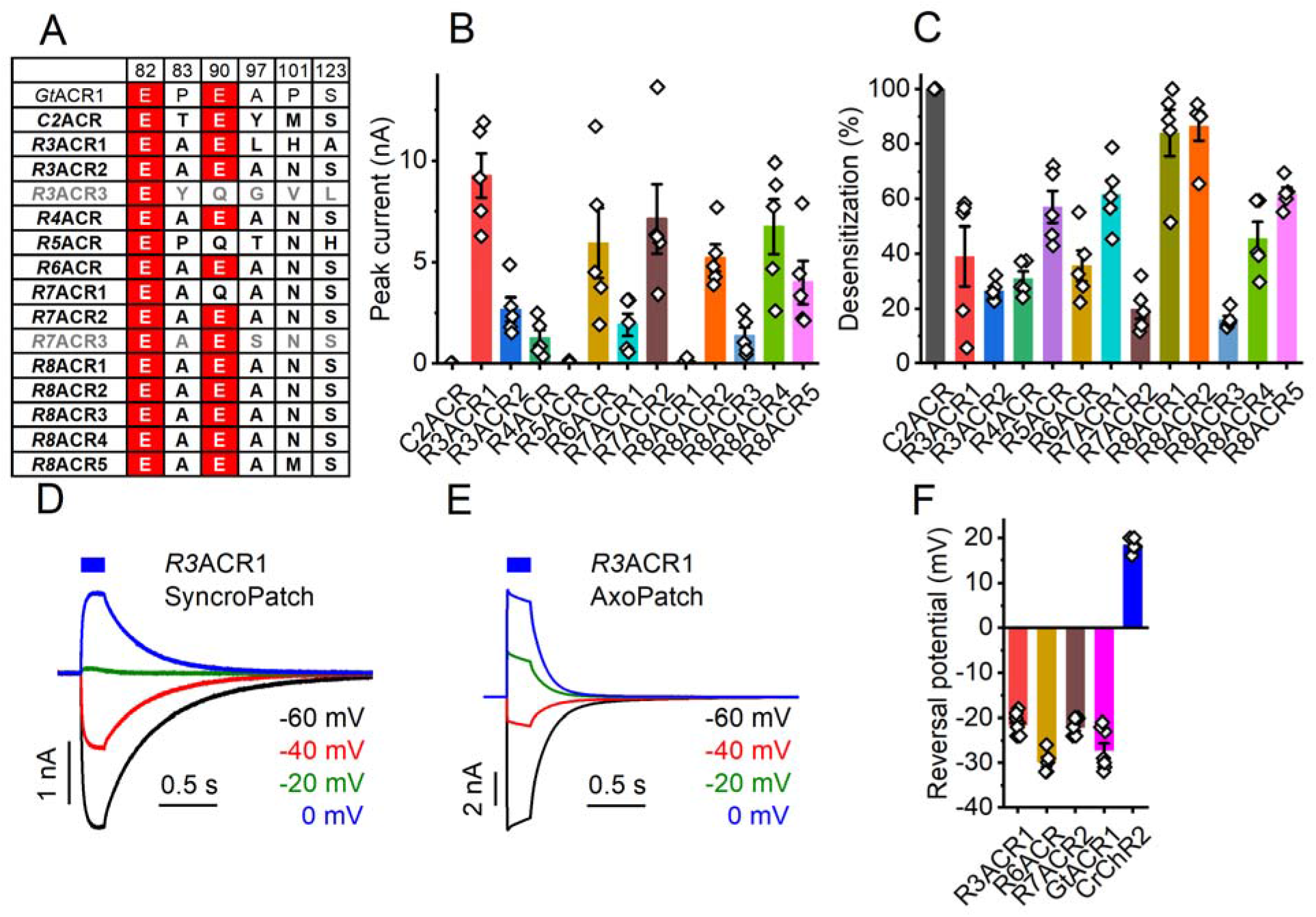
Cryptophyte ACRs. (A) Amino acid residues corresponding to the indicated positions in *Cr*ChR2. ChRs characterized in this study are in bold, non-functional homologs are in grey. Conserved glutamates are highlighted red. (B) Peak photocurrent amplitudes generated at −60 mV in response to 1-s light pulses at the wavelength of the spectral maximum. (C) Desensitization of photocurrents after 1-s illumination. (D and E) Series of photocurrent traces recorded from R3ACR1 upon incremental voltage increase with the SyncroPatch 384i (D) and AxoPatch 200B (E) at 470-nm excitation. Note the smaller amplitude and slower kinetics of the SyncroPatch traces, as expected from the lower stimulus intensity. (F) Reversal potentials of photocurrents. In B, C and F the bars show mean ± sem; empty diamonds, data from individual cells.

To verify permeability for anions in the three cryptophyte ACR homologs that were well-expressed and generated photocurrents in the nA range by manual patch clamp (Fig. 4B), we used a high-throughput automated patch clamp (APC) instrument, SyncroPatch 384i, with solutions provided by the manufacturer (for their full compositions see Table S1). The internal solution was predominantly CsF to promote formation of gigaseals, and the external solution was predominantly NaCl. Representative series of photocurrent traces recorded from *R3*ACR1 under incremental voltage using the SyncroPatch 384i and AxoPatch 200B with the same solutions are shown in Figs. 4D and E. *Gt*ACR1 and *Cr*ChR2, well-characterized by manual patch clamp, were included in the SyncroPatch experiment as ACR and CCR controls, respectively. With the SyncroPatch solutions, the E_rev_ of *Gt*ACR1 photocurrents was negative, whereas that of *Cr*ChR2 was positive (Fig. 4D). In all tested homologs the E_rev_ was close to that of *Gt*ACR1 (Fig. 4D), which confirmed their anion selectivity.

Previously, we and others demonstrated that Cysl33, Serl56 and Tyr207 in *Gt*ACR1 (absorption maximum 515 nm) corresponding to Argl29, Glyl52 and Phe203 in *Gt*ACR2 (absorption maximum 470 nm) define the spectral difference between these two proteins (15, 26)(E.G. Govorunova, O.A. Sineshchekov and J.L. Spudich, manuscript in preparation). According to *Gt*ACR1 crystal structures, the side chains of Cysl33 and Serl56 are located near the β-ionone ring of the chromophore (Fig. 5A), whereas the hydroxyl group of Tyr207 forms a hydrogen bond with Asp234 in the photoactive center. Comparative analysis of these positions (Fig. 5C) and action spectra of photocurrents (Fig. 5B and D) in the 13 functional ACR homologs has revealed that only those proteins in which the residues match those of *Gt*ACR2 exhibit blue-shifted absorption maxima. When Cys or Met are found at position 133, or Ser or Ala at position 156, the spectrum is shifted to longer wavelengths.

**Figure 5.**
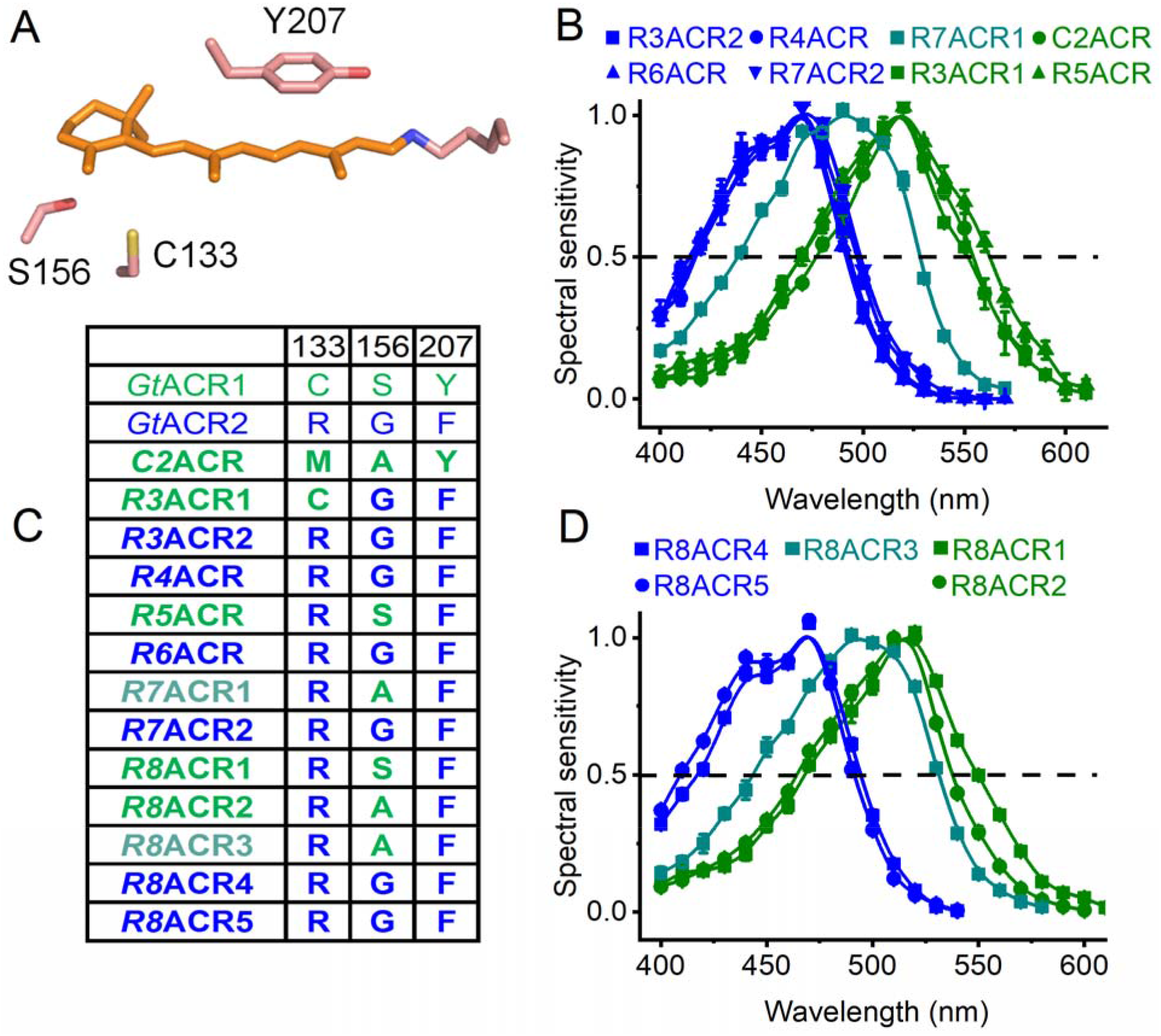
Color tuning in cryptophyte ACRs. (A) A crystal structure of *Gt*ACR1 (6EDQ) showing the three side chains that contribute to the spectral difference between *Gt*ACR1 and *Gt*ACR2. (B and D) Action spectra of photocurrents generated by the indicated cryptophyte ACRs. The data points show mean ± sem (n = 4-8 scans). (C) Amino acid residues involved in color tuning in the functional cryptophyte homologs. The numbering is according to the *Gt*ACR1 sequence.

### Dinoflagellate ChRs

Dinoflagellates exhibit genuine phototactic orientation (27–29), and their genomes encode multiple type I rhodopsins (30–32). However, to the best of our knowledge, none of these rhodopsins has so far been reported to exhibit channel function. Some rhodopsin sequences from dinoflagellates of the genera *Ansanella*, *Pelagodinum*, and *Symbiodinium* (6, 33-36) contain the TCP motif in the middle of TM3 that is conserved in most so far known channelrhodopsins (Fig. S3A). This motif is also conserved in 17 proteins encoded by the deep-ocean TARA marine transcriptomes that cluster together with these dinoflagellate rhodopsins and form a distinct branch of the phylogenetic tree (Fig. 1). A very unusual feature of this entire sequence cluster is that Asp212 of bacteriorhodopsin, highly conserved in all so far known channelrhodopsins, is replaced with Asn or, in one homolog, Leu (Fig. S3B).

Only one of the five tested metagenomic rhodopsin domains of this group was electrogenically photoactive upon expression in HEK293 cells, producing photocurrents barely resolved from the noise level (Fig. 6A, black bar). The fusion protein formed disk-shaped fluorescent aggregates within the cells (Fig. S3C, top). The addition of the trafficking signal (TS) between rhodopsin and EYFP, and the endoplasmic reticulum export motif (ER) at the C terminus of the fusion protein (37) reduced formation of the aggregates (Fig. S3C, bottom) and significantly increased the photocurrents, although they still reached only ~20-pA level at best (Fig. 6A, blue bar). In our standard buffer system with nearly symmetrical ionic concentrations in the bath and pipette the sign of the photocurrents reversed at positive voltages indicating passive ion transport (Fig. 6B, top). We named this protein mgdChR1 (for <u>m</u>etagenomic <u>di</u>noflagellate homolog <u>Ch</u>annel<u>R</u>hodopsin 1).

**Figure 6.**
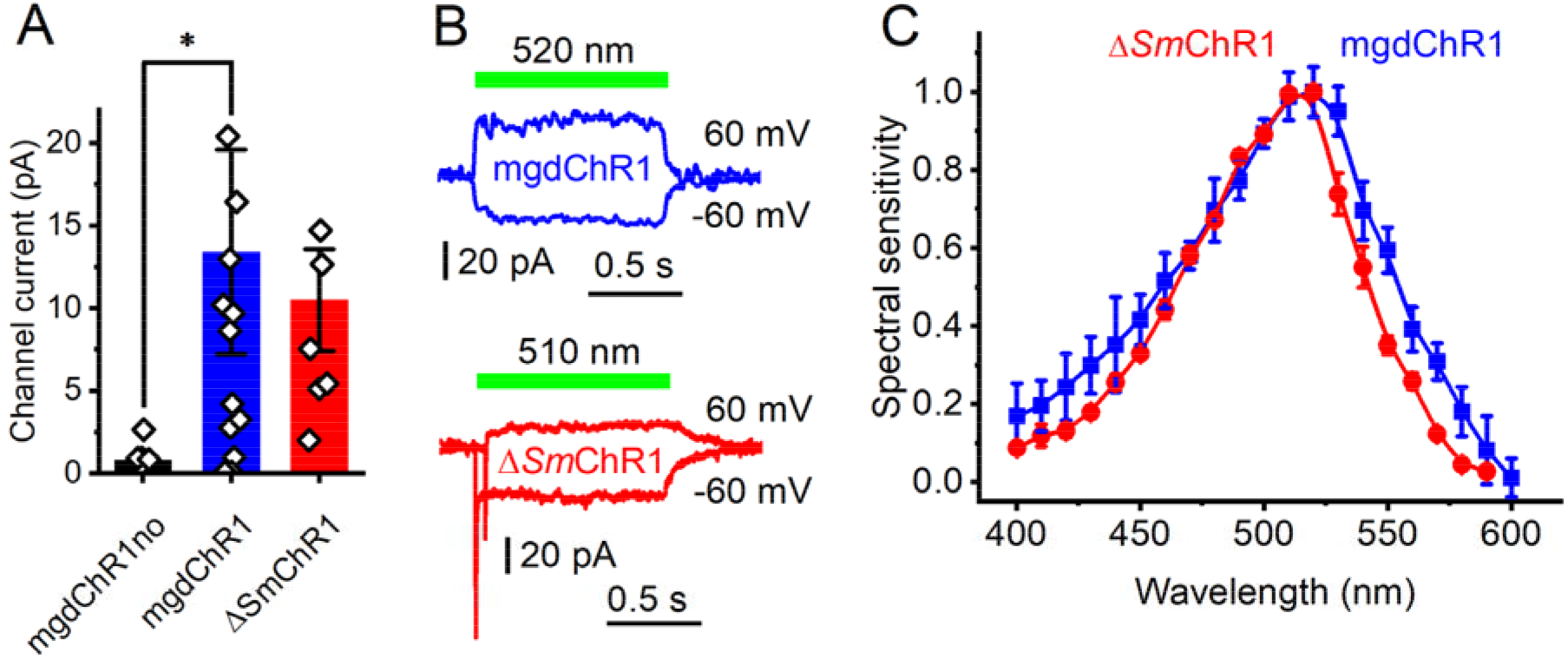
ΔSmChRl and its metagenomic homolog mgdChRl. (A) Peak photocurrent amplitudes generated at −60 mV in response to 1-s light pulses at the wavelength of the spectral maximum. The bars show mean ± sem; empty diamonds, data from individual cells. The asterisk indicates p< 0.01 by the Mann-Whitney test. “mgdChRlno” denotes the construct without TS and ER motifs. (B) Photocurrent traces recorded at −60 and 60 mV from mgdChRl (top) and ΔSmChRl (bottom). The ΔSmChRl at 60 mV was shifted 50 ms to the right relative to the trace at −60 mV to show the fast negative peak. (C) Action spectra of photocurrents. The data points show mean ± sem (n = 10-12 scans).

A homologous rhodopsin domain from the coral endosymbiont *Symbiodinium microadriaticum* has a ~300-residue N-terminal extension (Fig. S4), which is much longer than that found in other known ChRs, including mgdChR1. An expression construct encoding residues 1-600 produced no tag fluorescence. However, when the N-terminal extension was deleted, and TS and ER export motifs added, fluorescence was observed, and passive photocurrents of a similarly small amplitude as from mgdChR1 were recorded (Fig. 6B, bottom). We named this protein Δ*Sm*ChRl to emphasize truncation of the N-terminal extension. A homologous protein Δ*Sm*ChR2 from the same organism also generated photocurrents, but their amplitudes were even smaller. Channel currents from Δ*Sm*ChR1, but not from mgdChR1, were preceded with a fast negative current, the sign of which did not reverse at positive voltages (Fig. 6B, bottom). Such currents have previously been recorded from several other ChRs and interpreted as intramolecular charge displacement associated with isomerization of the retinal chromophore (24). The photocurrent action spectra of mgdChR1 and Δ*Sm*ChR1 peaked in the green spectral region (Fig. 6C). Small amplitudes of mgdChR1 and Δ*Sm*ChR1 photocurrents make accurate measurements of the reversal potentials problematic, so we were not able to determine their ionic selectivity.

## DISCUSSION

We report functional testing of 43 ChR homologs from prasinophytes, stramenopiles, cryptophytes, dinoflagellates and metagenomic samples. An unexpected result is that ACRs appear to be more widely spread among protist taxa than CCRs. Another unexpected result is that the same residue pattern comprising conserved Glu82 and Glu90 with non-carboxylate residues in the positions of Glu83, Glu97, Glu101, and Glu123 (*Cr*ChR2 numbering) found in most ACRs from stramenopiles, cryptophytes, haptophytes, and metagenomic samples is also found in the CCRs, *Cs*CCR and *Ms*CCR, from prasinophytes. Out of five Glu residues in TM2 and TM2-TM3 loop, Glu82 is most conserved across the entire ChR family. According to our empirical calculations using PROPKA3 (38), the pK_a_ of the Glu82 homolog is acidic in all so far published X-ray crystal structures of ChRs, including that of *Gt*ACR1 in which it apparently does not prevent anion conductance. In *Cr*ChR2, replacement of Glu82 with Ala strongly inhibited expression in mammalian cells judged by the tag fluorescence and correspondingly reduced photocurrents (39), which suggests that this residue is needed for correct protein folding and/or membrane targeting.

Glu90 appears to be essential for cation conductance in *Cr*ChR2, as mutation of this residue to Lys or Arg confers permeability for anions (19). Yet, this Glu is conserved in most ACRs except those from prasinophytes and Labyrinthulea. In the unphotolyzed state of both *Cr*ChR2 (40) and *Gt*ACR1 (41) this residue is neutral at neutral pH. Glu90 deprotonates during the photocycle of *Cr*ChR2 (42, 43). Photoinduced protonation changes of the Glu90 homolog in *Gt*ACR1 (Glu68) have not been studied by time-resolved molecular spectroscopy, but electrophysiological and UV-vis flash-photolysis data indicate that it also deprotonates upon photoexcitation (44). Further research is needed to clarify the role of this residue in anion conductance.

Photocurrent desensitization in different ChR families correlates with accumulation of different intermediates of the photocycle. In MerMAID1 desensitization is correlated with the M intermediate (22), but in *Rhodomonas* BCCRs it is with a novel extremely blue-shifted intermediate (23). Finally, desensitization in *Cr*ChR2 is correlated with accumulation of blue-absorbing P480 that is considered either as a late intermediate in a single branched photocycle (45) or the initial state of a parallel photocycle (43). Desensitization was reduced in the E44Q and C84T mutants of MerMAID1 (22). However, the mutated residues (corresponding, respectively, to Glu90 and Cys128 of *Cr*ChR2) are not the sole cause of strong desensitization in MerMAIDs, because they are conserved in many ChRs that do not show strong desensitization, including the closely related *s*TACR1 characterized here.

According to quantum mechanical/molecular mechanical calculations using the *Gt*ACR1 crystal structure, replacement of Ser156 with Gly or Ala stabilizes S_0_, predicting a 11-12-nm blue shift of the absorption maximum (46). All tested cryptophyte ACRs that contain Gly in this position exhibited blue-shifted spectra. The spectra of two ACRs that contain Ala in this position (R7ACR1 and R8ACR3) were ~25 nm blue-shifted from that of *Gt*ACR1, whereas the spectra of the other two (C2ACR and R8ACR2) were very similar to that of *Gt*ACR1, suggesting that the expected phenotypic effect of Ser to Ala substitution in these proteins was compensated for by other changed residues.

Our results and those of other groups suggest that most biophysical properties of ChRs relevant for their optogenetic applications cannot be assigned to a few individual residues, but rather reflect interactions between many of them. A cumulative larger set of electrophysiological data to which our study contributes might be used in the future to train machine learning algorithms to identify sequence motifs that define ionic selectivity, desensitization and absorption spectra. Implementation of such algorithms has already helped to improve plasma membrane targeting and light sensitivity of ChRs (47, 48).

Protein sequences of dinoflagellate ChRs and their metagenomic homologs are distantly related to ChRs from giant viruses (Fig. 1), two of which have been shown recently to passively conduct cations upon heterologous expression (7). However, Asp212 of bacteriorhodopsin is conserved in these viral CCRs, as in most other known microbial rhodopsins, whereas it is replaced with Asn in dinoflagellate ChRs. Analysis of *Symbiodinium* transcriptomes reveals potential latent infection by large dsDNA viruses (49), so viral origin of dinoflagellate ChRs cannot be excluded.

So far, the function of ChRs as photoreceptors guiding phototaxis has been verified directly only in the chlorophyte *C. reinhardtii*, the model organism for which methods of gene silencing and knockdown have been developed (1, 50). Several other chlorophyte and one cryptophyte species have been shown to generate photoreceptor currents, very similar to those in *C. reinhardtii* and likely resulting from ChR photoexcitation (51–54). The direction of photoreceptor currents recorded in both freshwater and marine flagellates is depolarizing, which reflects cation influx or anion efflux. Both *C. reinhardtii* phototaxis receptors are CCRs (2, 3), but ACRs might also contribute to depolarizing photoreceptor currents even in marine flagellates, if their membrane potential is sufficiently low. To the best of our knowledge, the membrane potential has not been estimated in any ACR-containing organism, but it is −170 mV in the giant marine unicellular alga *Acetabularia mediterranea* (55).

Based on the action spectra of dinoflagellate phototaxis, rhodopsins have been suggested as photoreceptors that guide this behavior (56). Our demonstration of channel activity in dinoflagellate rhodopsins with the TCP motif in TM3 strongly supports this hypothesis. The spectral sensitivity of dinoflagellate ChRs matches that of phototactic accumulation observed in *Symbiodinium* and unclassified coral symbiotic dinoflagellates (57, 58). The latter studies suggest that coral larvae use GFP fluorescence to attract dinoflagellate symbionts that are necessary for their survival.

Manual patch clamp is time-consuming and requires considerable skill. We sought to test whether APC can be used for characterization of hundreds of ChR variants that evolved in various protists. The planar-array principle implemented in the SyncroPatch 384i allows seal formation on micron-size orifices in the glass bottom of microwell plates (chips) into which cell suspension is pipetted, thus bypassing pipette fabrication and offering the option for recording multiple cells in parallel (59). APC is mostly used for drug screening, especially cardiac safety testing, in stably transfected cell lines. However, generation of such lines for ChR screening is not practical. We found that even upon chemical transfection that yielded only 30-70% visibly fluorescent cells depending on the construct, using the SyncroPatch 384i considerably sped up data collection, as compared to manual patch clamp.

## MATERIALS AND METHODS

### Bioinformatics

To identify metagenomic homologs of MerMAIDs, haptophyte ACRs and Labyrinthulea ACRs, we first searched selected datasets of the Integrated Microbial Genomes and Microbiomes at the Department of Energy’s Joint Genome Institute (JGI) (Data Set S4) using the keyword “rhodopsin”, and then performed blastp (protein-protein BLAST) search using RubyACR sequences as a query. A similar procedure was used to identify rhodopsin genes in the dinoflagellate genomes from various sources listed in Data Set S4. *Cafeteria roenbergensis* ChRs were identified in the National Center for Biotechnology Information (NCBI) non-redundant protein database using blastp and *Gt*ACR1 sequence as a query.

To explore the diversity of cryptophyte ACRs, we analyzed transcriptomes of 20 cryptophyte strains each sequenced on the illumina HiSeq 2000 platform and assembled with the Bridger algorithm (60). Using a hidden Markov model (HMM) (61) based on known cryptophyte ACRs, we identified 15 novel transcripts for experimental characterization. We also analyzed 136 deep-ocean metatranscriptomic libraries from the TARA Oceans Expedition (62) assembled with the Plass protein-level algorithm (63). Four distinct HMMs were built using previously known sequences of cryptophyte ACRs, cryptophyte BCCRs, chlorophyte CCRs and MerMAIDs. While many transcripts could be uniquely assigned to one of these four HMMs, some aligned weakly but equally well to two or more HMMs, and could not be assigned unambiguously. Remarkably, 17 of these “ambiguous” sequences turned out to be close homologs of dinoflagellate ChRs that were not included among our HMMs.

Rhodopsin sequences from Data Set S1 were aligned using MUSCLE with default parameters as implemented in MegAlign Pro software v. 17.1.1 (DNASTAR Lasergene, Madison, WI) and truncated after the end of TM7. Phylogeny was analyzed with IQ-TREE v. 2.1.2 (64) using automatic model selection and ultrafast bootstrap approximation (1000 replicates) (65). The best tree was visualized and annotated with iTOL v. 5.7 (66).

### Molecular biology and HEK293 transfection

DNA polynucleotides encoding the opsin domains optimized for human codon usage were synthesized and cloned by GenScript (Piscataway, NJ) into the mammalian expression vector pcDNA3.1 (Life Technologies, Grand Island, NY) in frame with an EYFP tag for expression in HEK293 cells. The cells were transfected using the ScreenFectA transfection reagent (Waco Chemicals USA, Richmond, VA). All-*trans*-retinal (Sigma) was added at the final concentration of 3 μM immediately after transfection.

### Manual patch clamp recording

Photocurrents were recorded 48-96 h after transfection in whole-cell voltage clamp mode with an AxoPatch 200B amplifier and digitized with a Digidata 1440A using pClamp 10 software (all from Molecular Devices, Union City, CA). Patch pipettes with resistances of 2-4 MΩ were fabricated from borosilicate glass. The ionic compositions of the bath and pipette solutions are shown in Table S1. For determination of E_rev_, K^+^ in the pipette solution was replaced with Na^+^ to minimize the number of ionic species in the system, and the holding voltages were corrected for liquid junction potentials calculated using the Clampex built-in calculator. Continuous light pulses were provided by a Polychrome V (T.I.L.L. Photonics GMBH, Grafelfing, Germany) in combination with a mechanical shutter (Uniblitz Model LS6, Vincent Associates, Rochester, NY; half-opening time 0.5 ms). The maximal photon density at the focal plane of the 40× objective was 5.2-8.5 mW mm^−2^ depending on the wavelength. The action spectra were constructed by calculation of the initial slope of photocurrent and corrected for the photon density measured at each wavelength (5). Further analysis was performed using Origin Pro software (OriginLab Corporation, Northampton, MA). The images were taken with a CoolSNAP HQ2 monochrome camera (Photometrics, Tucson, AZ).

### Automated patch clamp recording

Automated patch clamp recording was conducted with a SyncroPatch 384i (Nanion Technologies) using planar borosilicate glass medium-resistance chips in a 384 microtiter plate format with one or four holes per well and Nanion Standard solutions (for their composition see Table S1). Transfected cells were dissociated using TrypLE^TM^ Express, diluted with CHO-S-SFM-II medium (both from ThermoFisher) and resuspended in External Physiological solution (Nanion Technologies) at 10^5^ −4×l0^5^ cells ml^−1^. Each well was filled with 30 μl Chip Fill solution, to which 20 μl of the cell suspension was added. Seal formation was enhanced by the addition of 40 μl of NMDG 60 solution with 10 mM CaCl_2_ (final concentration). After capturing the cells, 50 μl of the external solution was replaced with 40 μl of NMDG 60 solution, and 40 μl of the mixture was removed. For data acquisition and analysis, respectively, PatchControl384 and DataControl384 software v. 1.9.0 was used (both Nanion Technologies). Illumination was provided with LUXEON Z blue LEDs LXZ1-PB01 (470 ± 20 nm) controlled by custom-built software.

### Statistics

Descriptive statistics was used as implemented in Origin software. The data are presented as mean ± sem values; the data from individual replicates are also shown when appropriate. The sample size was estimated from previous experience and published work on similar subjects, as recommended by the NIH guidelines. No normal distribution of the data was assumed; when a specific statistics hypothesis was tested, non-parametric tests were used.

## Data availability

The polynucleotide sequences of ChR homologs reported in this study have been deposited to GenBank (accession numbers MW557552-MW557594).

## Acknowledgements

We thank Dr. Tim Strassmaier, Leo Angelo Morada, Stephan Holzhauser and Rasmus Gönner (Nanion Technologies) for their expert help with the SyncroPatch 384i.

## Funding

This work was supported by National Institutes of Health Grants R01GM027750 and U01NS118288, and Endowed Chair AU-0009 from the Robert A. Welch Foundation to J.L.S, and by the Natural Sciences and Engineering Research Council of Canada (NSERC) Discovery Grant RGPIN-2018-04397 to L.S.B. A.P. was supported by a NSERC USRA scholarship, S.C. was supported by the Agricultural Science and Technology Innovation Program (ASTIP). The access to the SyncroPatch 384i was provided by a research grant from Nanion Technologies. The work conducted by the U.S. Department of Energy Joint Genome Institute, a DOE Office of Science User Facility, is supported by the Office of Science of the U.S. Department of Energy under Contract No. DE-AC02-05CH11231. The content is solely the responsibility of the authors and does not necessarily represent the official views of the National Institutes of Health.

## Conflict of interest

The authors declare no conflict of interest.

## Supplemental Results

### “Core” chlorophyte and streptophyte ChR homologs

Only one Glu residue (corresponding to Glu82 of CrChR2) is conserved in TM2 of the functionally characterized *Pyramimonas* ACRs (6). Four sequences from Chlorophyceae (two from *Chlamydomonas noctigama*, one from *Chlamydomonas sp*. and one from Chloromonas subdivisa) also exhibit this residue pattern (Fig. S1B). Upon expression of three of these polynucleotides, small hyperpolarizing photocurrents that did not reverse at positive voltages were recorded (Fig. S1C). They likely reflect intramolecular transfer of the Schiff base proton to an outwardly located acceptor, as previously found in other ChRs (24, 67). The *C. subdivisa* sequence was very poorly expressed and generated no currents.

Out of the four Chlorodendrophyceae sequences with the misplaced Asp212 homolog, one (named here *Tch*ChR) has already been tested earlier and found to be non-electrogenic (68). We have synthesized and tested two other sequences of this group, PsChR4 from *P. subcordiformis* and *Ta*ChR from *T. astigmatica*. Neither generated photocurrents upon expression in HEK293 cells, although normal tag fluorescence was observed.

Functional CCRs have previously been reported in the streptophyte classes Mesostigmatophyceae (53) and Klebsormidiophyceae (69). A ChR homolog has also been identified in *Coleochaete irregularis* (6) from the class Coleochaetophyceae, which is more closely related to land plants. However, we could not detect any photocurrents upon its expression in HEK293 cells.

## Supplemental Figure Legends

**Figure S1.**
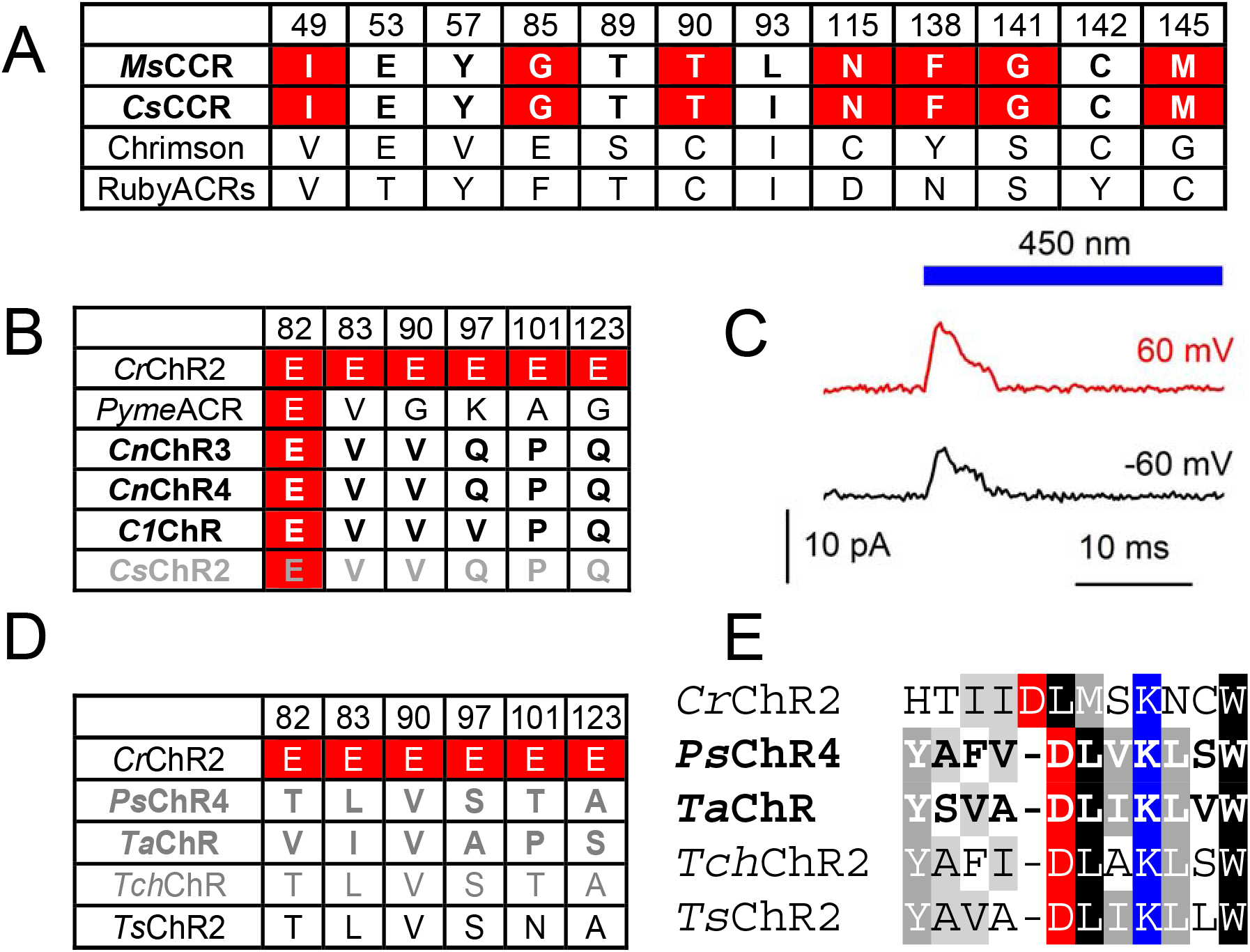
(A) The residues of the retinal-binding pockets. Variants tested in this study are in bold. The residues in *Ms*CCR and *Cs*CCR that differ from those in Chrimson and RubyACRs, earlier known red-shifted ChRs, are highlighted red. The numbers are according to bacteriorhodopsin sequence. (B and D) The residues in the positions of the conserved glutamates (highlighted red) in the ion conductance pathway in the indicated homologs from Chlorophyceae (B) and Chlorodendrophyceae (D). Variants tested in this study are in bold, non-functional, in gray. The numbers are according to *Cr*ChR2 sequence. (C) Photocurrent traces recorded from *C1*ChR in response to 1-s illumination at −60 and 60 mV. (E) Alignment of the part of TM7 of the indicated Chlorodendrophyceae homologs. The Schiff base Lys is highlighted blue; the upstream Glu, red.

**Figure S2.**
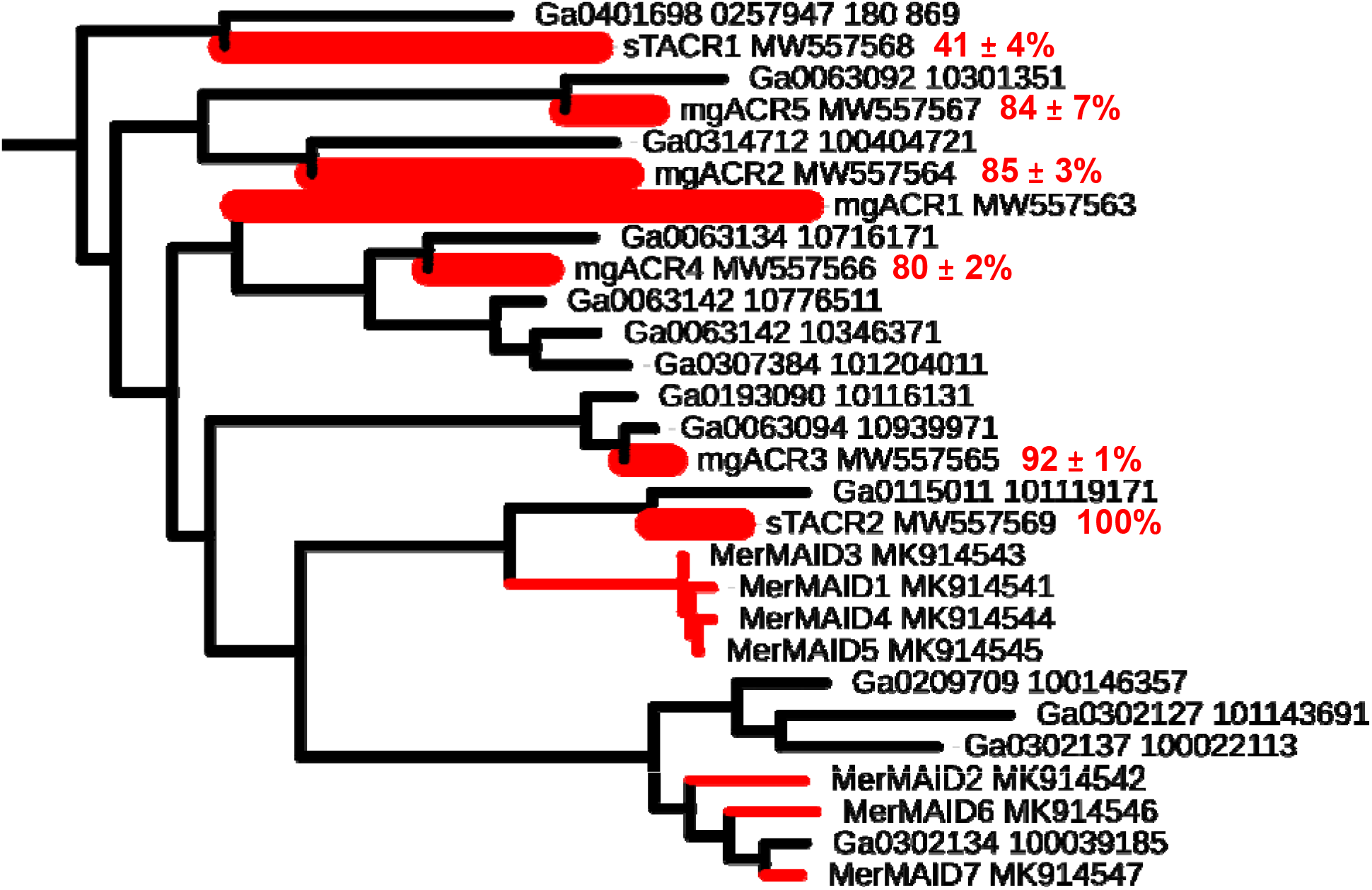
A section of the phylogenetic tree from Fig. 1 redrawn in a rectangular format. Red nodes show variants with proven anion selectivity, thicker nodes show variants tested in this study. Red numbers are the values of photocurrent desensitization from Fig. 3C in the main text (mean ± sem, n = 5-6 cells).

**Figure S3.**
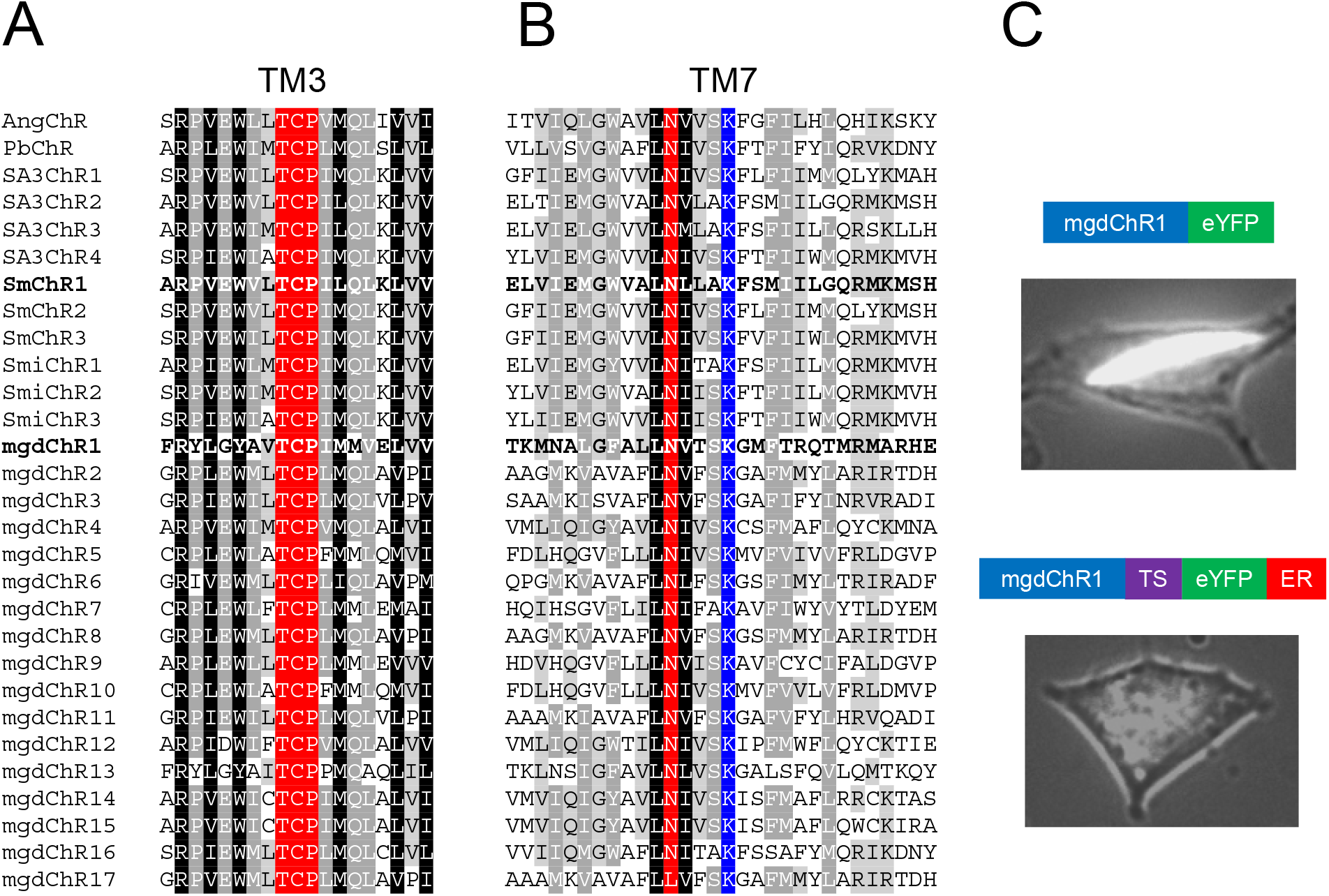
(A and B) Alignments of TM3 (A) and TM7 (B) of dinoflagellate ChRs and their metagenomic homologs. The TCP motif and the Asn residue corresponding to Asp212 of bacteriorhodopsin are highlighted red. (C) EYFP tag fluorescence in cells expressing mgdChR1 constructs schematically shown on top of the images. TS, trafficking signal, ER, endoplasmic reticulum export motif.

**Figure S4.**
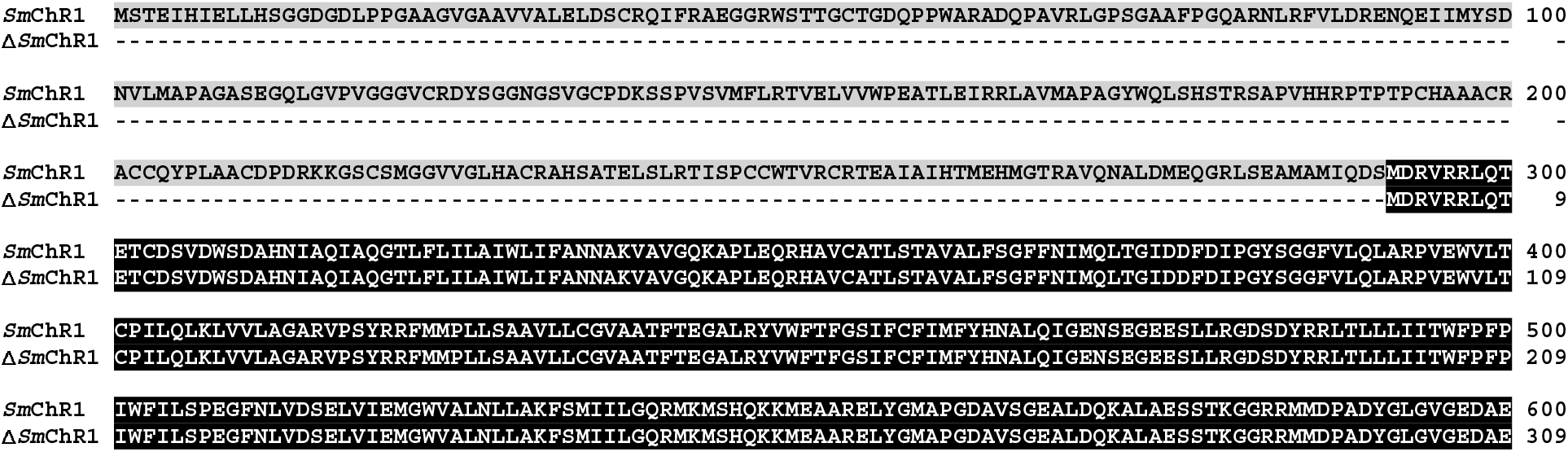
Alignment of *S. microadriaticum* ChR1 with and without the N-terminal extension. Supplemental Table Legends

## Supplemental Table Legends

**Table S1.** Compositions of the solutions used in patch clamp recording. Abbreviations: Asp, aspartate; EGTA, ethylene glycol tetraacetic acid; HEPES, 4-(2-hydroxyethyl)-1-piperazineethanesulfonic acid; UP, liquid junction potential; NMDG, N-Methyl-D-glucamine. All concentrations are in mM.

|  | NaCl | KCl | CsCl | CsF | CaCl<sub>2</sub> | MgCl<sub>2</sub> | EGTA | HEPES | NMDG* | Glucose | NaAsp | pH | LJP<br>mV |
| --- | --- | --- | --- | --- | --- | --- | --- | --- | --- | --- | --- | --- | --- |
| **Pipette standard** | — | 126 |  |  | 0.5 | 2 | 5 | 25 | — | — | — | 7.4 | — |
| **Pipette NaCl** | 126 | — |  |  | 0.5 | 2 | 5 | 25 | — | — | — | 7.4 | — |
| **Bath standard** | 150 | — |  |  | 1.8 | 1 | — | 10 | — | 5 | — | 7.4 | 1.1 |
| **Bath NaAsp** | — | — |  |  | 1.8 | 1 | — | 10 | — | 5 | 150 | 7.4 | -13 |
| **Bath pH 6.4** | 150 | — |  |  | 1.8 | 1 | — | 10 | — | 5 | — | 6.4 | 1.1 |
| **Bath NMDG** | 1.4 | — |  |  | 1.8 | 1 | — | 10 | 148.6 | 5 | — | 7.4 | 7 |
| **Nanion Internal CsF** | 10 | — | 10 | 110 | — | — | 10 | 10 | — | — | — | 7.2 | — |
| **Nanion External Chip Fill** | 140 | 4 | — | — | — | — | — | 10 | — | 5 | — | 7.4 | — |
| **Nanion External Physiological** | 140 | 4 | — | — | 2 | 1 | — | 10 | — | 5 | — | 7.4 | — |
| **Nanion External NMDG 60** | 80 | 4 | — | — | 2 | 1 | — | 10 | 60 | 5 | — | 7.4 | — |

\*NMDG stock (pH 9) was prepared from 1M NMDG and ~700 mM HCl.

**Table S2.** List of cryptophyte strains analyzed. Abbreviations: BEA, Banco Español de Algas, of the Universidad of Las Palmas de Gran Canaria, Spain; CCAC, Culture Collection of Algae at the University of Cologne, Germany; SCCAP, Scandinavian Culture Collection of Algae and Protozoa at the University of Copenhagen, Denmark; CCMP, Culture Collection of Marine Phytoplankton at the Provasoli-Guillard National Center for Marine Algae and Microbiota at Woods Hole Oceanographic Institution, USA; NIES, National Institute for Environmental Studies, Tsukuba, Japan

|  | Catalog number | Genus and species names | Habitat (growth medium) | ACRs found? |
| --- | --- | --- | --- | --- |
| 1. | CCMP 0268 | *Chroomonas* sp. | Marine (ASP-12)* | No<sup>#</sup> |
| 2. | CCAC 4037 B | *Chroomonas* sp. | Marine (ASP-12)* | No<sup>#</sup> |
| 3. | NIES 1370 | *Chroomonas* sp. | Marine (ASP-12)* | No<sup>#</sup> |
| 4. | CCAC 0173 B | *Chroomonas* sp. | Marine (ASP-H)* | No<sup>#</sup> |
| 5. | CCAC 1627 B | *Chroomonas* sp. | Marine (ASP-H)* | Yes |
| 6. | CCAC 3782 B | *Chroomonas* sp. | Freshwater | No<sup>#</sup> |
| 7. | CCAC 3670 B | *Chroomonas* sp. | Marine (ASP-H)* | No<sup>#</sup> |
| 8. | BEA 0199B | *Cryptomonas* cf. *pyrenoidifera* | Freshwater | No<sup>#</sup> |
| 9. | CCAC 0108 | *Cryptomonas gyropyrenoidosa* | Freshwater | No<sup>#</sup> |
| 10. | CCAC 0031 | *Cryptomonas obovoidea* | Freshwater | No<sup>#</sup> |
| 11. | CCAC 0064 | *Cryptomonas ovata* | Freshwater | No<sup>#</sup> |
| 12. | SCCAP K0416 | *Geminigera* sp. | Marine (ASP-12)* | No<sup>#</sup> |
| 13. | CCAC 1074 B | *Komma caudata* | Freshwater | No<sup>#</sup> |
| 14. | CCMP 0760 | *Rhodomonas* sp. | Marine (ASP-12)* | Yes |
| 15. | CCAP 979/6 | *Rhodomonas* sp. | Marine (ASP-12)* | Yes |
| 16. | CCAC 1480 B | *Rhodomonas* sp. | Freshwater | Yes |
| 17. | CCAC 3787 B | *Rhodomonas* sp. | Marine (ASP-12)* | Yes |
| 18. | CCAC 3407 B | *Rhodomonas* sp. | Marine (ASP-12)* | Yes |
| 19. | CCAC 3799 B | *Rhodomonas* sp. | Marine (ASP-12)* | Yes |
| 20. | BEA 0603B | *Urgorri complanatus* | Marine (ASP-12)* | No<sup>#</sup> |

\*For media recipes see the CCAC website (https://www.uni-due.de/biology/ccac/)
<sup>#</sup>Our finding of no ACR transcripts in this species does not necessarily mean that they are not encoded by its genome and expressed; the transcript abundance may simply have been below the detection limit of our method.

## Supplemental Data Set Legends

### Data Set S1.

Genbank accession numbers, abbreviated protein names, source organisms, habitats, transcript names and amino acid sequences used to construct the phylogenetic tree in Fig. 1. Note that only sequences that cover the entire N-terminal and rhodopsin domains are included. Literature references to identification and electrophysiological characterization of the sequences are also provided. The sequences identified and characterized in this study are shown in bold.

### Data Set S2.

A Newick file of the tree shown in Fig. 1 in the main text.

### Data Set S3.

An alignment of the C-truncated sequences used to construct the tree in Fig. 1 in the main text.

### Data Set S4.

A list of sequence databases searched.

## Notes

### Competing Interest Statement

The authors have declared no competing interest.

